# Using data-dependent and independent hybrid acquisitions for fast liquid chromatography-based untargeted lipidomics

**DOI:** 10.1101/2023.10.12.562117

**Authors:** Kanako Tokiyoshi, Yuki Matsuzawa, Mikiko Takahashi, Hiroaki Takeda, Mayu Hasegawa, Junki Miyamoto, Hiroshi Tsugawa

**Affiliations:** Department of Biotechnology and Life Science, Tokyo University of Agriculture and Technology, 2-24-16 Nakamachi, Koganei-shi, Tokyo 184-8588, Japan; RIKEN Center for Sustainable Resource Science, 1-7-22 Suehiro-cho, Tsurumi-ku, Yokohama, Kanagawa 230-0045, Japan; RIKEN Center for Brain Science, 2-1 Hirosawa, Wako, Saitama 351-0106, Japan; Department of Applied Biological Science, Tokyo University of Agriculture and Technology, Fuchu-shi, Tokyo 183-8509, Japan; RIKEN Center for Integrative Medical Sciences, 1-7-22 Suehiro-cho, Tsurumi-ku, Yokohama, Kanagawa 230-0045, Japan; Molecular and Cellular Epigenetics Laboratory, Graduate School of Medical Life Science, Yokohama City University, Tsurumi-ku, Yokohama, Kanagawa 230-0045, Japan

## Abstract

Untargeted lipidomics using liquid chromatography coupled with tandem mass spectrometry (LC-MS/MS) has become an essential technique for large cohort studies. When a fast LC gradient of less than 10 min is used for the rapid screening of lipids, the annotation rate decreases because of the lower coverage of the MS/MS spectra caused by the narrow peak width. We propose a systematic procedure to achieve a high annotation rate in fast LC-based untargeted lipidomics by integrating data-dependent acquisition (DDA), and sequential window acquisition of all theoretical mass spectra data-independent acquisition (SWATH-DIA) techniques with the updated MS-DIAL program. Our strategy uses variable SWATH-DIA methods for quality control (QC) samples, which are a mixture of biological samples analyzed multiple times to correct MS signal drifts. In contrast, biological samples are analyzed using DDA to facilitate the structural elucidation of lipids using the pure spectrum to the maximum extent. We demonstrate our workflow using an 8.6 min LC gradient, where QCs are analyzed using five different SWATH-DIA methods. The results indicated that using both DDA and SWATH-DIA achieves 2.0-fold annotation coverage from publicly available benchmark data obtained by a fast LC-DDA-MS technique and offers 94.5% lipid coverage compared with the benchmark dataset from a 25 min LC gradient. Our study demonstrated that harmonized improvements in the analytical conditions and informatics tools provide a comprehensive lipidome in fast LC-based untargeted lipidomics, not only for large-scale studies but also for small-scale experiments, contributing to both clinical applications and basic biology.

---

Lipid metabolism is an important and complex process that provides essential components of cellular membranes, signaling molecules, and energy resources for living organisms. Untargeted lipidomics using liquid chromatography coupled with tandem mass spectrometry (LC-MS/MS) has become an essential technique for understanding the lipidome profile of biospecimens in which 500–1000 unique molecules can be characterized by the MS/MS spectrum in combination with retention time (RT) behaviors^1^, leading to an understanding of lipid diversity and its biological importance. Several studies have acquired large-scale LC-MS/MS data to create a lipidome atlas by analyzing multiple organs and/or biological species and mapping lipidome cartography encompassing the spatiotemporal alteration of a lipid molecule.^2–4^ In addition, untargeted lipidomics has been used in population-scale human cohort studies, providing biologically and clinically significant discoveries and validating potential biomarkers, such as plasma ceramide profiling for cardiovascular disease.^5–7^

Experimental designs that acquire more than 1,000 samples require a framework for high-throughput LC-MS measurements. A high-throughput measurement system minimizes the data size and measurement errors within a batch and reduces data acquisition times, thereby improving the accuracy of peak annotation and alignment procedures with the small RT tolerance. Faster measurement methods using short LC gradients or ion mobility spectrometry-mass spectrometry have been developed in several studies.^3, 8, 9^ In contrast, the short LC method provides less coverage for MS/MS acquisition, which is the most important information source for lipid annotation, resulting in lower annotation rates compared to a conventional LC gradient of 20–30 min. To elucidate the structure of lipids, data-dependent acquisition (DDA) is the gold standard technique in lipidomics because the MS/MS spectrum contains few contaminants owing to a strict isolation window for the precursor *m/z*. However, the narrow peak width in fast LC reduces the MS/MS acquisition rates in a scan cycle, resulting in inadequate MS/MS assignment of the precursor ions using DDA techniques (**Figure 1a**).

**Figure 1.**
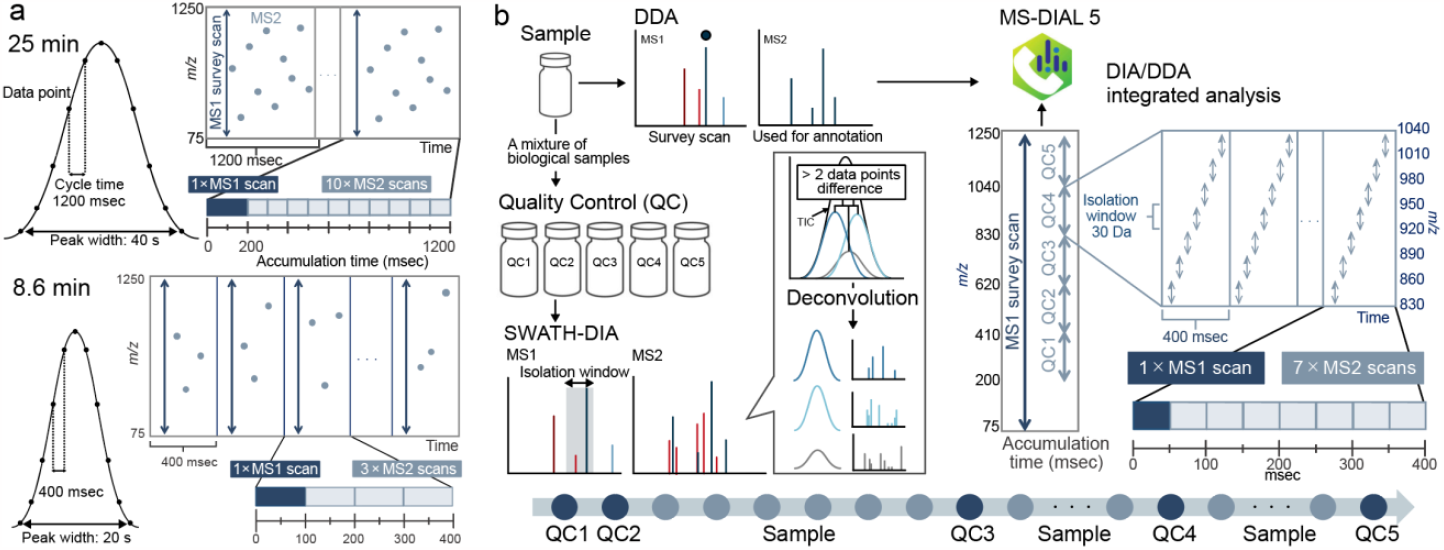
Overview of hybrid MS-based lipidomics. (a) Differences of peak width and example settings for data-dependent acquisition (DDA) in 25 and 8.6 min LCs. Two LC gradient settings that have already been reported were used in this study. The cycle time and DDA conditions were determined to acquire approximately 15 data points for a single peak on average. (b) Summary of experimental design using quality control (QC) samples. The approach combines both sequential window acquisition of all theoretical mass spectra data-independent acquisition (SWATH-DIA) and DDA for QC measurements and samples, respectively. Five QC measurements are used in a batch sequence, each of which has a different setting for precursor isolations. The isolation window was fixed to 30, and the 210 Da precursor range was scanned in a QC analysis, covering the mass range of 200–1250 Da for MS/MS acquisitions that grasp major lipid molecules. The deconvolution algorithm distinguishes co-eluted peaks if the peak top difference exceeds two data points: the cycle time is set to 0.4 while the average peak width is calculated as 20 s in fast LC. The updated MS-DIAL was used to integrate the hybrid MS data.

In contrast to DDA, data-independent acquisition (DIA) acquires MS/MS data for all ions in principle, and it has garnered attention for the untargeted analysis of biomolecules.^10^ Still, few studies have investigated the effective use of both DIA and DDA for fast LC-MS measurements, and the development of analytical chemistry and related informatics techniques is an emerging need in studies using lipidomics.

We believe that the structural elucidation of unknown lipids should be performed mostly using a pure and less contaminated MS/MS spectrum, which is achieved by DDA and targeted product-ion scans. In contrast, the comprehensive annotation of known lipids can be accelerated by DIA because a comprehensive library template of the precursor *m/z* and its theoretical fragment pattern of molecules can be prepared in lipidomics, allowing the identification of diagnostic ions in a contaminated spectrum. In this study, we propose a novel strategy using both DIA and DDA in an analytical batch sequence to increase annotation rates in fast LC-MS, where a single LC-MS measurement is set to take 8.6 min for the showcase. To analyze the complex data structure effectively, we developed an algorithm to integrate DIA and DDA data simultaneously, which was implemented in MS-DIAL.^1^ The program also accepts variable sequential window acquisition of all theoretical mass spectra (SWATH)-DIA under different experimental conditions. We integrated the DDA and DIA data from five different SWATH-DIA measurements. Our procedure was evaluated using the National Institute of Standards and Technology (NIST) SRM 1950 plasma and mouse feces, which are important biopsies for clinical applications.

## EXPERIMENTAL SECTION

### Chemicals and Reagents

The ammonium acetate solution for high-performance liquid chromatography (HPLC), ethylenediaminetetraacetic acid (EDTA), acetonitrile (MeCN), methanol (MeOH), 2-propanol (IPA), and ultrapure water (H_2_O) of quadrupole time-of-flight MS (QTOF-MS) grade were purchased from FUJIFILM Wako Pure Chemical Corp. (Osaka, Japan). Methyl tert-butyl ether (MTBE) for HPLC was purchased from Sigma-Aldrich (Tokyo, Japan). The SRM 1950 plasma was purchased from NIST (USA).

### Fecal sample

Animal experiments were conducted in accordance with the ethical protocol approved by the Tokyo University of Agriculture and Technology (R5-50). C57BL/6J male mice were purchased from SLC (Shizuoka, Japan). The mice were fed the chow of CE-2 (CLEA Japan, Tokyo, Japan). Mouse feces were harvested, frozen immediately, and stored at −80 °C until lipid extraction.

### Lipid extraction

Lipid extraction was performed on ice according to a previous study using a biphasic solvent system comprising cold MeOH, MTBE, and H_2_O.^11^ See **Supplementary Information** for details of the extraction procedure. As a summary, lipids from 20 μL and 5 mg were resolved in MeOH whose solvent was stored in LC-MS vials. In this study, the same material was used for QC and biological samples for system evaluations. The sample was analyzed 20 times by DDA and analyzed five times by the SWATH-DIA methods described below. Only one injection was performed from a LC-MS vial.

### Fast LC-MS measurement

LC conditions were established according to a previous study^3^ in which the original system was slightly modified for the analytical column, solvent, and hardware conditions used. The LC system comprised a Nexera X2 UPLC system (Shimadzu, Kyoto, Japan). Lipids were separated on an Imtakt UK-C18 MF column (50 × 2 mm:3 μm) (Imtakt, Kyoto, Japan). The MS detection of lipids was performed using QTOF-MS (ZenoTOF 7600; SCIEX, Framingham, MA, USA). The DDA method was set in both positive and negative ion modes as follows: MS1 and MS2 mass ranges, *m/z* 75–1250; MS1 accumulation time, 200 ms; Q1 resolution, units; MS2 accumulation time, 100 ms; maximum candidate ions, 3. The SWATH-DIA method was set in both positive and negative ion modes as follows: MS1 accumulation time, 100 ms; MS2 accumulation time, 50 ms; Q1 window, 30 Da; and MS1 mass range, *m/z* 75–1250. Five different SWATH-DIA settings were prepared in which the precursor scanning range for MS/MS acquisition was set to 300–510, 510–720, 720–930, 930–1140, and 1140–1250 Da. In the negative-ion mode, the scanning ranges for MS/MS were set to 200–410, 410–620, 620–830, 830–1040, and 1040–1250 Da. See **Supplementary Information** for the other details of LC-MS conditions. The 30 Da isolation window was determined based on the annotation rate when analyzing fecal samples where the annotation count of lipids was evaluated in the SWATH-DIA experiment, covering the precursor ions at *m/z* 600–810. A total of five isolation window settings of 10, 20, 30, 40, and 50 Da were investigated, where the cycle time was fixed at approximately 500 ms and the MS2 accumulation time was changed accordingly. The annotation results are summarized in **Supplementary Table 1**.

### Untargeted lipidomics data using a conventional LC and a similar fast LC setting as a benchmark

The raw data from a conventional LC-MS measurement and a similar fast LC-MS measurement (25 and 6.8 min for a single run, respectively) were downloaded from the RIKEN DROP Met website (http://prime.psc.riken.jp/menta.cgi/prime/drop_index) under the DM0031 index. The data utilized was obtained from “1 Srm1950 Human Plasma Lipidomics”^1^. Among these, untargeted analysis using SCIEX 6600-based DDA measurements for four NIST SRM plasma replicates provided the highest annotation rate, covering 495 unique lipid molecules, and was used as a benchmark in this study. Additionally, an untargeted analysis using Thermo Q-Exactive Plus-based DDA measurements for three NIST SRM plasma replicates was used as previously reported results of fast LC-MS measurement.

### MS-DIAL development and analysis

The MS-DIAL program was updated to efficiently integrate and analyze the DDA and DIA data simultaneously. The updated program recognized the precursor window setting directly from wiff/wiff2 format data. In addition, users can independently set the data acquisition type, that is, DDA, SWATH, or all-ion fragmentation, for each analysis file. For the SWATH-DIA data, an MS/MS chromatogram deconvolution algorithm was performed according to a previous study^12^. Individually acquired MS/MS spectral outputs and annotation results were integrated into the alignment results. In this study, biological samples were analyzed using the DDA mode, whereas QC samples were analyzed using SWATH-DIA in which five QCs covered the spectra for the entire scanning range. The MS/MS spectrum with the highest annotation score was assigned as the representative spectrum for each aligned spot. Thus, the spectrum from DDA is often assigned to the alignment result, even if a QC sample contains spectral information for the same peak spot. SWATH-DIA covers the MS/MS spectrum which was not acquired by DDA. The algorithm was implemented in the MS-DIAL program. The RT library, which was prepared using a machine learning method in a previous study for 25 min LC-MS measurement, was also updated. This method uses the LC gradient curve information. First, the RT in the original database optimized for 25 min LC was converted to the solvent B% value of the LC gradient. Second, the time at the B% value in the 8.6 min LC gradient was used for the lipid RT. The data processing parameters are listed in **Supplementary Table 2**. The annotated results were curated manually. The representative adduct forms used to calculate the annotation rates are listed in **Supplementary Table 3**. The lipidome results are available in **Supplementary Data 1**, where one fecal sample was excluded because of a large mass error caused by missed calibration in the SCIEX calibration delivery system.

## RESULTS AND DISCUSSION

### Overview of fast LC-based lipidomics using DDA and SWATH-DIA hybrid acquisitions

Our procedure followed the guidelines proposed by the metabolomics/lipidomics standards initiatives, which recommend the use of QC samples to monitor the sensitivity drifts of MS in inter- and intra-batches (**Figure 1b**)^13^. We propose that QCs be measured using SWATH-DIA to acquire comprehensive MS/MS spectra, while biological samples be measured using DDA. Although MS/MS chromatogram deconvolution can reconstruct a pure spectrum for each precursor peak from co-eluted metabolites, the spectrum becomes artificial, resulting in difficulty in the elucidation of novel lipids that have previously been unknown. In contrast, MS/MS spectrum is essential for lipid structure description because only the precursor *m/z* value provides many candidates for lipid isomers. Thus, our procedure facilitates the comprehensive profiling of known lipids using both SWATH-DIA and DDA while the spectrum of DDA in the alignment results is used to the maximum extent to elucidate unknown peaks (see Experimental section).

We described three important conditions for comprehensive lipid annotation in fast LC-MS measurements. First, the isolation window of SWATH-DIA was set to less than 30 Da to increase the annotation rates with the deconvolution algorithm, because the larger isolation width caused lower annotation rates despite the larger accumulation times in our investigation (**Supplementary Table 1)**. Although a precursor separation window of 30 Da provided the best annotation rate in SWATH-DIA, the optimal window depended on the LC gradient and *m/z* scanning ranges of interest. Second, precursor scanning using SWATH-DIAs should cover the *m/z* range whose MS/MS spectra are essential for annotation at the molecular species level (e.g., PC 16:0_18:2 instead of PC 36:2) in a batch sequence: PC, 16:0, 18:2, and 36:2 denote phosphatidylcholine, a fatty acid (FA) moiety containing 16 carbons, an FA moiety containing 18 carbons and two unsaturated degrees, and the summed value of carbons and unsaturated degrees from the two acyl chains, respectively. Finally, the cycle time was optimized to acquire 10–20 data points in a chromatographic peak, which is the minimum requirement for quantification in analytical chemistry. In this study, we used five SWATH-DIA settings, each of which covered the ∼210 Da range of the precursor *m/z* with an isolation window of 30 Da, indicating that QC samples were measured at least five times to obtain MS/MS spectra in the 200–1250 Da precursor ion range. Analyzing QCs five times in a single analytical batch is practical; two pre- and post-QC measurements (four in total) are usually performed to monitor MS sensitivity and RT stability before and after an analytical sequence. Thus, an additional measurement of QC in an intra-batch setting is the minimum requirement for this analysis. This implies that even with a small experimental design of 10–20 samples, comprehensive MS/MS acquisition can be achieved with a fast LC technique. As the number of samples increases the analyses of QCs, the precursor isolation window per QC may become narrower in a large-scale lipidomic study, providing more purified spectra for lipid annotation. The accumulation times for MS1 and MS2 in SWATH-DIA were set to 100 and 50 ms, respectively, to detect low-abundance diagnostic ions^12^. The dataset was analyzed using the updated MS-DIAL program.

### Validation of untargeted lipidomics using fast LC-MS measurements

We prepared a simple dataset to evaluate hybrid MS techniques using a standard plasma material, NIST SRM 1950. The plasma was analyzed 20 times by DDA coupled with the 8.6 min LC method, and the same samples were analyzed by five different SWATH-DIA methods, two of which were analyzed before the DDA analyses, one of which was analyzed after 10 injections by DDA, and the others were analyzed after the DDA analyses. We also downloaded untargeted lipidomic data of the same plasma sample using DDA coupled with a 25 min LC gradient as a benchmark, whose annotation rate outperformed those of other methods in a previous study. We excluded duplicate annotations from different ion forms, for instance, ceramides could be assigned as [M + H]^+^, [M − H]^−^, and [M + CH3COO]^−^, and the number of unique annotated molecules was compared among the experiments.

Higher annotation rates in fast LC-MS measurements were obtained by integrating the DDA and SWATH-DIA data (**Figure 2a**). The number of annotated lipids in the DDA measurement was 268, whereas a set of data acquired using SWATH-DIA through five injections provided 350 annotated lipids. This is because of the lower coverage of MS/MS acquisitions using a single DDA file. We confirmed this hypothesis by increasing the amount of DDA data in the MS-DIAL analysis, where the annotation rate increased with an increase in imported DDA data. The annotation rate in analyzing the DDA datasets exceeded that of a single set of SWATH-DIAs when seven DDA datasets were analyzed simultaneously. In addition, the simultaneous analysis of 20 DDAs and SWATH-DIAs yielded 468 unique lipid molecules, whereas the analysis of 20 DDAs alone resulted in 403 lipids. This result can be interpreted as the MS/MS spectrum of extremely low-abundance ions obtained only in SWATH-DIA, whereas the contaminant spectrum of SWATH-DIA, which was not purified by the deconvolution algorithm, was compensated for by the spectrum of DDA in the alignment process. We validated this hypothesis using the following two examples: The MS/MS spectrum of a ganglioside subclass, GM3, whose intensity was extremely low in plasma samples, was not obtained in the DDA analysis, whereas the spectra were obtained using SWATH-DIA **(Figure 2b**). In contrast, the MS/MS spectrum of ether-linked lysophosphatidylcholine (LPC O-) containing 18:1 (LPC O-18:1, RT 2.0 min, *m/z* 508.3767) was annotated in the DDA measurement (**Figure 2c**) while it was not characterized in SWATH-DIA data (**Figure 2d**) owing to the interference of highly abundant MS/MS chromatographic peaks of lysophosphatidylcholine (LPC) including 16:0 (LPC 16:0, RT 1.9 min, *m/z* 496.3403). The product ions of *m/z* 184 and 104 whose existences in MS/MS are the minimum requirement to define the LPC O-subclass were not recognized as the product ion spectrum of LPC O-18:1 in the deconvolution process (**Figure 2e**). These results demonstrated the advantages of using both DDA and SWATH-DIAs. Although the annotation rate did not reach 495, which was obtained from publicly available 25 min LC-MS measurements, our fast LC-MS data acquisition technique provided a 1.98-fold annotation rate when compared to the result from a fast LC-MS technique (**Supplementary Table 4**). Similar findings were observed in the mouse fecal samples (**Figure 2f**). The increased coverage of the fecal lipidome would lead to the discovery of new bio-active bacteria-dependent lipids, known as *N*-acyl amides and glycosyl ceramides, whose bioactivity is confirmed at extremely low concentrations of the nanomolar order.^14^ Because plasma and feces are popular biopsies in clinical applications, our fundamental investigation using a fast LC-MS technique will contribute to future large-scale cohort studies. Details on the annotation count are provided in **Supplementary Data 2**.

**Figure 2.**
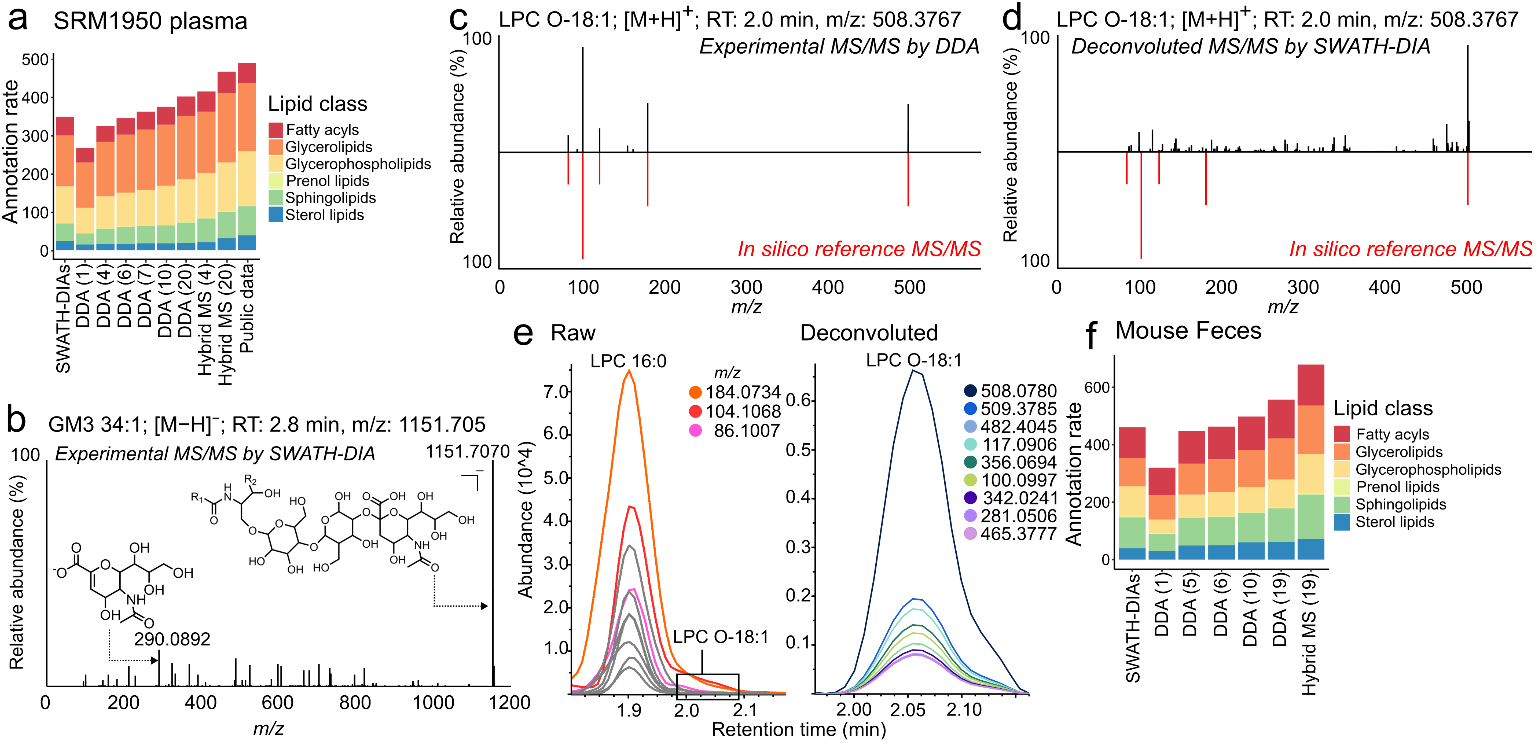
Increase of annotation rates by hybrid MS approach. (a) Annotation results for SRM 1950 plasma. The X-axis represents the analytical method while the Y-axis shows the annotation rate. “SWATH-DIAs” denotes the results from analyzing the dataset acquired using five SWATH-DIA experiments. “DDA (1–20)” represents the results from analyzing 1–20 sets of data using DDA. The annotation count in the DDA data analysis exceeds the results from the SWATH-DIAs by more than seven. “Hybrid MS (4) or (20)” correspond to the integrated analysis of the SWATH-DIA sequence and 4 or 20 DDA analysis data, respectively. “Public data” signifies the results obtained from the publicly available data containing four replicates of NIST plasma acquired by DDA with a 25 min gradient LC method. (b) MS/MS spectrum annotated as GM3 34:1;O2 obtained by the SWATH-DIA analysis. (c) MS/MS spectrum of ether-linked lysophosphatidylcholine containing 18:1 (LPC O-18:1, RT 2.0 min, *m/z* 508.3780) in DDA. (d) Deconvoluted MS/MS spectrum of LPC O-18:1 in the SWATH-DIA measurement. (e) MS/MS chromatograms where the product ions are derived from the precursor ions of 479–510 Da range. (f) Annotation results for fecal samples.

## CONCLUSION

We proposed a procedure to achieve high annotation rates in fast LC-MS measurements utilizing both SWATH-DIA and DDA techniques, which was showcased by the lipidomics of human plasma and mouse fecal samples. This methodology offers several key advantages, including (1) flexibility for both small- and large-scale studies, (2) scalability to large cohort studies owing to QC measurements for MS drift corrections, and (3) coverage of MS/MS spectra for comprehensive annotation of known lipid molecules and for structure elucidation of unknown MS/MS spectra. The procedure could not be achieved without updating the MS-DIAL algorithm that can integrate the DDA and variable SWATH-DIA-derived data. Considering the versatility, reproducibility, and potential for elucidating the diversity of lipidomes in living organisms, we believe that this approach becomes the gold standard for untargeted lipidomics in various studies, for understanding lipid metabolism linked to biological phenotypes, and for discovering new lipid structures that have been previously unknown.

## Supporting information

Source data for figures

## ASSOCIATED CONTENT

### Supporting Information

The Supporting Information is available free of charge on the ACS Publications website. The detail of lipid extraction and the fast LC-MS/MS setting. (**Supplementary Information**) (Microsoft Word Document) The annotation results of analyzing fecal samples to determine the isolation window. (**Table S1**); MS-DIAL parameters. (**Table S2**); Detail of Adduct form used for the calculation of the annotation rate. (**Table S3**); Annotation results from several LC-MS/MS measurements. (**Table S4**); Lipidomic results of biological samples. (**Supplementary Data 1**); Details of the annotation counts by lipid class for each analysis. (**Supplementary Data 2**) (Microsoft Excel Workbook)

### Data availability

All raw MS data are available on the RIKEN DROPMet website (http://prime.psc.riken.jp/menta.cgi/prime/drop_index) under the index number DM0050. The source data for the figures were also recorded as **Source Data**. The source code of MS-DIAL is available at https://github.com/systemsomicslab/MsdialWorkbench.

## AUTHOR INFORMATION

### Author Contributions

Hiroshi Tsugawa designed the study. Hiroshi Tsugawa and Y.M. updated the MS-DIAL program. K.T. and Hiroaki Takeda optimized the LC-MS conditions. M.H. and J.M. conducted the mouse experiments. M.T. updated the Lipid Spectral Database. K.T. performed the data analysis. K.T. and Hiroshi Tsugawa wrote the manuscript. All authors have thoroughly discussed this project and helped improve the manuscript.

## ACKNOWLEDGMENT

This study was supported by the JSPS KAKENHI (21K18216 to H.T.), the National Cancer Center Research and Development Fund (2020-A-9, H.T.), AMED Japan Program for Infectious Diseases Research and Infrastructure (21wm0325036h0001, H.T.), AMED Brain/MINDS (JP15dm0207001 to H.T. and H.T.), JST National Bioscience Database Center (NBDC, H.T.), and JST ERATO “Arita Lipidome Atlas Project” (JPMJER2101, H.T.).

---

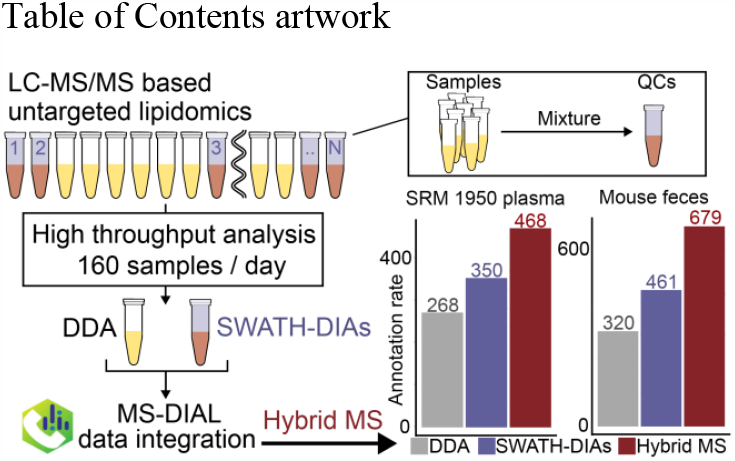

