## Supplementary Information for "Using data-dependent and independent hybrid acquisitions for fast liquid chromatography-based untargeted lipidomics"

**Corresponding author**

**Contents**

Detail of lipid extraction

Detail of fast liquid chromatography coupled with tandem mass spectrometry setting

Supplementary Table 1-4

Supplementary Data 1 and 2

Source data

**Lipid extraction**

Lipid extraction was performed on ice using a biphasic solvent system comprising cold MeOH, MTBE, and H_2_O.^1^ NIST SRM 1950 plasma was dispensed into 1.5 mL tubes of 20 µL each, and a total of ten tubes were used in this study. 20 μL of plasma sample was ultrasonicated with 225 μL MeOH for 10 min. After adding 750 μL of MTBE, vortexed for 10 s, shaken for 10 min, and centrifuged at 16,000 rpm for 3 min. Then, 800 μL of supernatant was collected in a clean tube, and mixed with 164 μL of H_2_O by a vortex mixer for 10 s. After centrifuging at 16,000 rpm for 3 min, 350 μL of the upper organic phase collected in a clean tube was dried up by centrifugal evaporator and resuspended in 100 μL MeOH. The solvents from 10 tubes were collected in a single clean tube and vortexed for 10 s. After centrifuging at 16,000 rpm for 3 min, the supernatant was transferred into LC-MS vials (Agilent Technologies, Santa Clara, California, USA) of 20 μL each. Only one injection was administered in the LC-MS vial. Fecal samples were homogenized using a multi-bead shocker with a metal cone (YASUI KIKAI, Japan) at 2500 rpm for 15 s, and 1 mL of MeOH was added. MeOH containing 5 mg of fecal sample was transferred to a new 2.0 mL tube, and additional MeOH was added to bring the total volume to 225 μL. The remaining procedure was the same as that used for plasma samples. In this study, 20 samples were analyzed by DDA, five samples were analyzed by the SWATH-DIA sequence described below, and the same material was used for quality control (QC) and biological samples for evaluation.

**Fast LC-MS measurements**

LC conditions were established according to a previous study^2^ in which the original system was slightly modified for the analytical column, solvent, and hardware condi-tions used. The LC system comprised a Nexera X2 UPLC system (Shimadzu, Kyoto, Japan). Lipids were separated on an Imtakt UK-C18 MF column (50 × 2 mm:3 µm) (Imtakt, Kyoto, Japan). Throughout the fast LC-MS measurement (8.6 min), the column was maintained at 65 °C and a flow rate of 0.6 mL/min. The mobile phases comprised (A) 1:1:3 (v/v/v) MeCN:MeOH:H_2_O with ammonium acetate (5 mM) and 10 nM EDTA and (B) 9:1(v/v) IPA:MeCN with ammonium acetate (5 mM) and 10 nM EDTA. A sample volume of 1 μL was used for the injection. The separation was performed under the following gradient: 0 min 0.1% (B), 0.1 min 0.1% (B), 0.2 min 15% (B), 1.1 min 30% (B), 1.4 min 48% (B), 5.6 min 82% (B), 6.9 min 99% (B), 7.1 min 99% (B), 7.2 min 0.1% (B), and 8.6 min 0.1% (B). The temperature of the rack was maintained at 4 °C.

The MS detection of lipids was performed using QTOF-MS (ZenoTOF 7600; SCIEX, Framingham, MA, USA). The DDA method was set in both positive and negative ion modes as follows: MS1 and MS2 mass ranges, *m/z* 75–1250; MS1 accumulation time, 200 ms; Q1 resolution, units; MS2 accumulation time, 100 ms; maximum candidate ions, 3; and CAD gas, 7. The following settings were used for positive/negative ion mode, independently: ion source gas 1, 40/50 psi; ion source gas 2, 80/50 psi; curtain gas, 30/35 psi; source temperature, 250/300 °C; spray voltage, 5500/-4500 V; declustering potential, 80/-80 V; and collision energy, 40/-42 ± 15 eV. The SWATH-DIA method was set in both positive and negative ion modes as follows: MS1 accumulation time, 100 ms; MS2 accumulation time, 50 ms; Q1 window, 30 Da; and MS1 mass range, *m/z* 75 – 1250. The other settings were the same as those used for the DDA. Five different SWATH-DIA settings were prepared in which the precursor scanning range for MS/MS acquisition was set to 300–510, 510–720, 720–930, 930–1140, and 1140–1250 Da. In the negative-ion mode, the settings were slightly changed when the scanning ranges for MS/MS were set to 200–410, 410–620, 620–830, 830–1040, and 1040–1250 Da. The 30 Da isolation window was determined based on the annotation rate when analyzing fecal samples. The annotation count of lipids was evaluated in the SWATH-DIA experiment, covering the precursor ions at *m/z* 600–810. A total of five isolation window settings of 10, 20, 30, 40, and 50 Da were investigated, where the cycle time was fixed at approximately 500 ms and the MS2 accumulation time was changed accordingly.

**Supplementary Tables**

Table S1. MS methods and the annotation results of analyzing fecal samples to determine the isolation window.

Table S2. MS-DIAL parameters.

Table S3. Detail of Adduct form used for the calculation of the annotation rate.

Table S4. Annotation results from several LC-MS measurements.

**Supplementary Data**

Supplementary Data 1. Lipidomic results of biological samples.

Supplementary Data 2. Details of the annotation counts by lipid class for each analysis.

**Source Data**

Source Data contains the source information for all figures.
